# Comparative phylogeography of the freshwater mussels of the southeastern United States: reconstructing historic drainages using molecular data

**DOI:** 10.1101/2021.03.08.434420

**Authors:** Scott T. Small, John P. Wares

## Abstract

Knowledge of species ages and their distribution enhance our understanding of processes that create and maintain species diversity at both local and regional levels. The largest family of freshwater mussels (Unionidae), reach their highest species diversity in drainages of the southeastern united states. By sequencing multiple loci from mussel species distributed throughout the drainages in this region, we attempt to uncover historical patterns of divergence and determine the role of vicariance events on the species formation in mussels and extend our hypothesis to freshwater animals in general. We analyzed 346 sequences from five genera encompassing 37 species. Species were sampled across 12 distinct drainages ending either in the Atlantic Ocean or the Gulf of Mexico. Overall the topologies of the different genera returned phylogenetic trees that were congruent with geographically contiguous drainages. The most common pattern was the grouping between the Atlantic slope and gulf coast drainages, however the Tennessee drainage was often the exception to this pattern grouping with the Atlantic slope. Most mussel species find a most recent common ancestor within a drainage before finding an ancestor between drainages. This supports the hypothesis of allopatric divergence followed by later burst of speciation within a drainage. Our estimated divergence times for the Atlantic-Gulf split agree with other studies estimating vicariance in fish species of the Atlantic and gulf coast.

---

The goal of comparative phylogeography is to evaluate the topological and chronological congruence of molecular phylogenies across a common biogeographic barrier in multiple species pairs (Avise 1998; Edwards and Beerli 2000). Comparisons among species allow inference into common mechanisms that have influenced the evolutionary, demographic, and distributional histories of taxa in an ecological region (Bermingham and Moritz 1998). Knowledge concerning the different ages of species and their areas of extent serve to enhance our understanding of processes that create and maintain species diversity at both local and regional levels. Improving our understanding of processes that promote and maintain diversity will allow us to focus conservation efforts on biologically diverse regions containing endemic species.

The southeastern United States (U.S.) provides the perfect example of endemic regional biodiversity as it contains high levels of endemic freshwater species such as gastropods, fish, crayfish, and mussels (Master 1990). There have been two proposed hypotheses to explain contemporary species distribution and diversity in the southeastern U.S, *i)* source-dispersal and *ii)* multiple vicariance. Under the hypothesis of source-dispersal, it is believed that a single ancestral drainage was a cradle for most of the present species diversity; contemporary patterns of species distribution are then due to coastal flooding, headwater captures, and anadromous fish migration which allow dispersal of species among rivers (Sepkoski and Rex 1974). Under the hypothesis of vicariance biogeography, a once widespread ancestor species was fragmented into smaller isolated populations by the formation of a barrier to gene flow; these species then diverged becoming reproductively isolated (Pflieger 1971; Mayden 1985; Mayden 1987; Wiley and Mayden 1985). These two hypotheses are by no means mutually exclusive and attempts to treat them as such may lead to data outliers that cannot be explained by either mechanism. A better hypothesis would incorporate mechanisms of both vicariance and source-dispersal to explain contemporary pattern of species diversity and distribution.

Despite the presence of mussels throughout the southeastern U.S., it has not been tested whether the distribution of freshwater mussels can be explained by either of these hypotheses. In this paper we use freshwater mussel taxa that are distributed throughout the drainages of the southeastern U.S. to test for both topological and chronological congruent patterns of biogeographic vicariance and dispersal. Freshwater mussels are ideal for this task because of an extrinsic link to fish dispersal; mussels do not migrate except during their larval life stage where they rely on fish in a parasitic life cycle. Hence any movement made by fish during times of sea level change or head-water capture would be preserved in the genome of freshwater mussels. By sequencing multiple loci from mussel species distributed throughout the drainages of the southeastern U.S., we can uncover historical patterns of divergence and determine the role of vicariance events on the species formation in mussels and freshwater animals in general.

If we find that multiple mussel taxa support a single divergence time it would follow that a single dispersal/vicariant event was most likely responsible for species formation, linking species in evolutionary history. Lieberman (2000) advocated the importance of identifying community-wide events as they serve to link species within an evolutionary framework and create a stable community (Lieberman 2000). Species in a stable community share an evolutionary history important to both the biotic stability of the community and inference of environmental factors affecting both past and future diversity (Lieberman 2000). Incorporating history of the entire community and its implied stability allow us to draw inferences on the long-term sustainability of species relationships within that community, an important factor for conservation management.

Here we use three mitochondrial loci (*16S ribosomal RNA, cytochrome c oxidase subunit I (CO1), NADH dehydrogenase subunit 1 (NAD1))* in a comparative phylogeographic approach to determine the relative importance of glacial and interglacial cycles on speciation of freshwater mussels. We test the congruence of divergence times among species to determine the stability of contemporary mussel communities. We focused on five widespread genera (*Alasmidonta, Elliptio, Lampsilis, Pleurobema*, and *Villosa*) that were sampled from the Altamaha (GA), Satilla (GA), Apalachicola-Flint-Chattahoochee (GA, AL, FL), Alabama-Coosa (GA, AL), Savannah (GA), Tennessee (TN), Ocklockonee (FL), Ogeechee (GA), St. Mary’s (FL) St. John’s (FL), and Choctowhatchee (FL). We utilized Bayesian phylogenetic construction and incorporate genealogical variance to estimate possible divergence times as well as the likelihood of multiple vicariance events. We use data from five mussel genera to answer the following questions: how old is the MRCA for each species group? Are molecular phylogenies similar in topology and chronology? Is there evidence of a phylogeographic break across the Appalachian Divide?

## Materials and Methods

We targeted 12 drainage basins covering an area from the Mississippi River east to south of the Pee Dee River, encompassing a majority of the rivers in the southeastern U.S. (Figure 1). We grouped drainages into either the Atlantic Slope or the eastern Gulf of Mexico Coastal Plains (Gulf Coast) based on the terminus of the rivers. The Atlantic Slope contains the Altamaha, Savannah, Satilla, Ogeechee, and Santee rivers and the Gulf of Mexico contains the Apalachicola-Chattahoochee-Flint (ACF), Coosa, Mobile, Choctawhatchee, Suwannee, Ochlockonee, and Tennessee. The Appalachian Divide separates the Atlantic Slope region and the eastern Gulf Coast, effectively isolating the Unionid fauna of the coastal rivers from direct contact with the species of the interior basin (Johnson 1970). The rivers and their freshwater faunas constitute discrete biogeographic systems (Sepkoski and Rex 1974).

**Figure 1.**
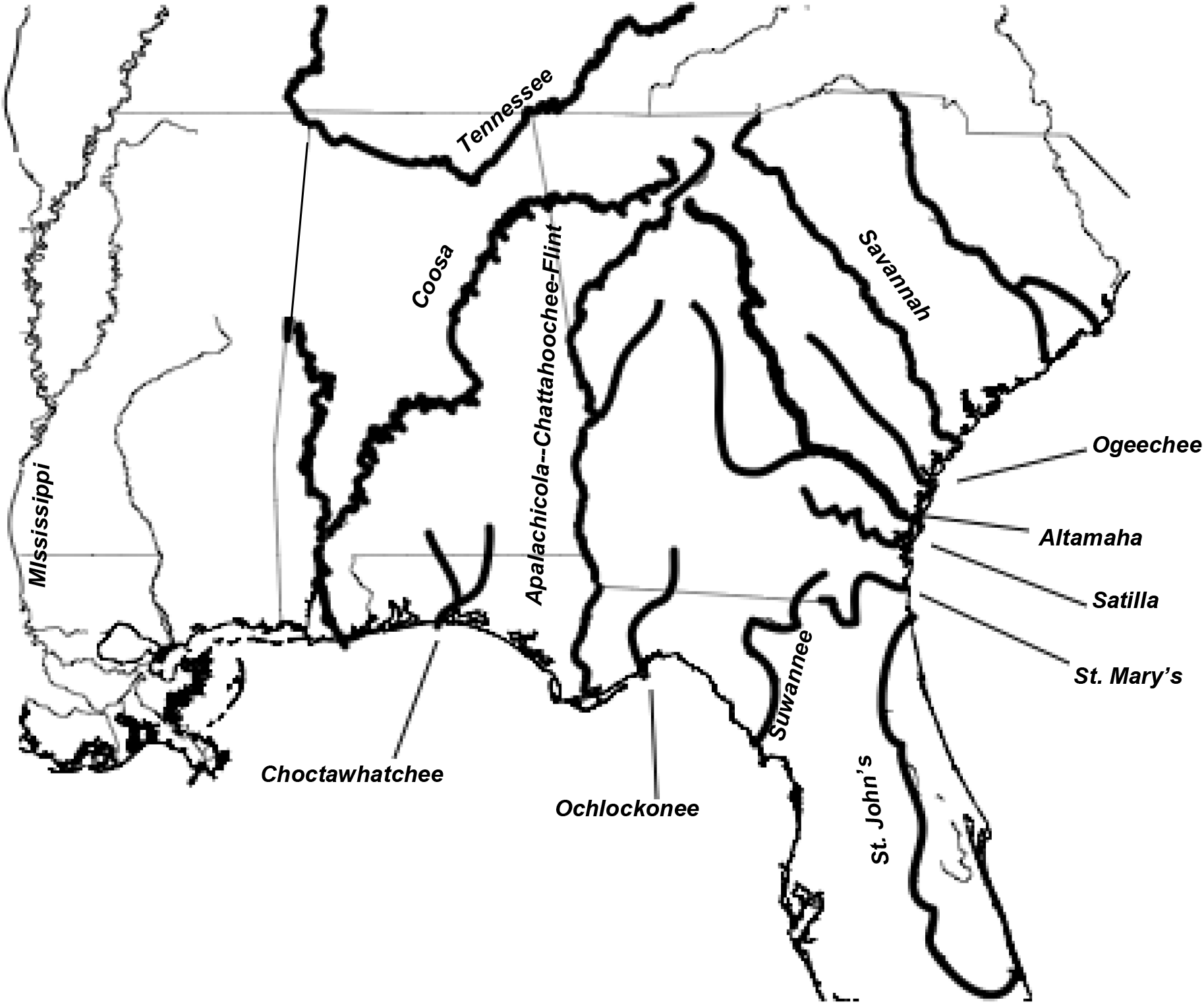
Map of southeast drainages. A map of the southeast United States showing in bold the rivers sampled for this project. The Mississippi is labeled for reference only and was not included in this study.

Mussel specimens were collected between 2007-2009 from taxonomic collections at Auburn University (AU), University of Alabama (UA), and North Carolina State (NC) (Supplementary Table S1). We focused our specimen collection on five genera, *Alasmidonta, Elliptio, Lampsilis, Pleurobema, Villosa*, that co-occur in most of the drainages (Table 1). From each genus we avoided species under taxonomic revision (Williams and Bogan 2008), species with purported anthropogenic introductions, and widespread congeners consisting of subspecies.

**Table 1.**
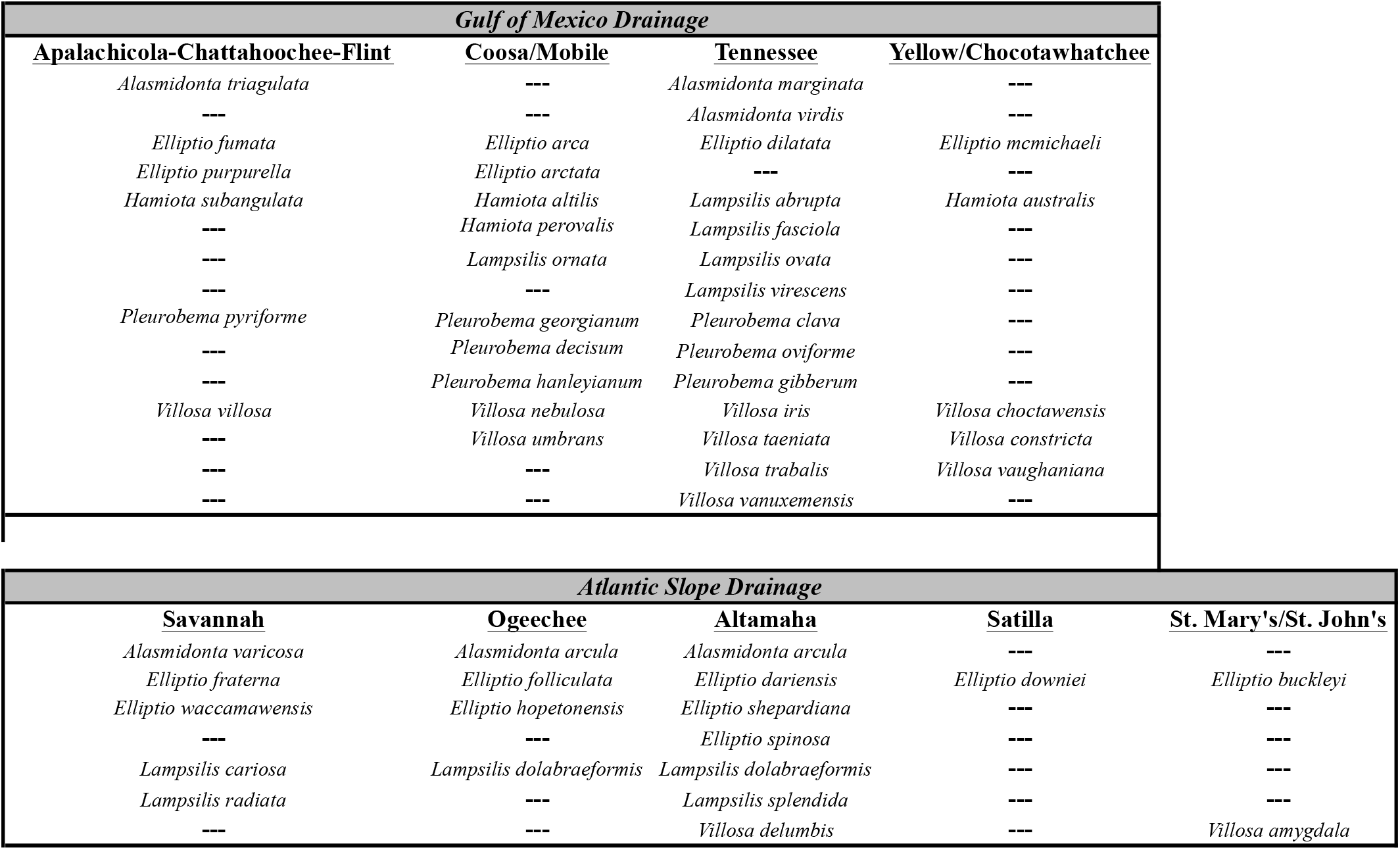
Mussel specimens collected from drainages on the Gulf Coast and Atlantic Slope.

Tissue was collected from the foot, abductor mussel, or mantle of mussel specimens and preserved in 95% EtOH for later DNA isolation. DNA from the collected tissue was then isolated using a modified CTAB (cetyl trimethylammonium bromide) isolation protocol based on Campbell, Serb et al. (2005). Samples were first homogenized using liquid nitrogen and micro-pestles. We next added CTAB to a final volume of 300ul with an addition of 25mg/ml Proteinase K solution (Gentra). Samples were allowed to digest at 55° C for 1-2 hours or until no solid tissue remained in the tube. DNA was precipitated using a chloroform wash step followed by the addition of 100% isopropanol. DNA was then eluted in 40ul of H_2_O and stored at -80° C until time of use.

We used PCR to amplify two loci (*16S, CO1*) from the mitochondrial genome. For mitochondrial loci (*16S, CO1*) we obtained primer sequences from Campbell, Serb et al. (2005) and Folmer, Black et al. (1994), respectively. Additional *16S, CO1*, and *NAD1* sequences were obtained from Genbank (queried January 2009; Supplementary Table 2). PCR amplifications were performed in 20µl volumes consisting of 0.5μM each primer, 0.8mM total dNTPs, 3mM MgCl_2_, and 1U Taq polymerase (Promega). Annealing temperatures for each locus were as follows: *16S*, 50° and COI, 40°. PCR products were prepared for sequencing using Exonuclease I and Antarctic Phosphatase (New England Biolabs, Ipswich, MA, USA). Sequencing reactions were carried out in 10µl volumes with 80ng of prepared template, 0.6μM primer, 0.6μl BigDye Terminator (Applied Biosystems, Foster City, CA, USA) and 3.4 μl Better Buffer (The Gel Company). Sequence reactions were cleaned and precipitated with 4 volumes 75% isopropanol, suspended in Hi-Di formamide (Applied Biosystems, Foster City, CA, USA) and run on an ABI 3730 at the University of Georgia. For each locus, sequence data were edited using CodonCode Aligner v.2.06 (CodonCode Corporation, Dedham, MA, USA). Sequences were aligned using Aligner’ sbuilt-in ‘end-to-end’ algorithm, examined and edited for likely artifacts caused by poly-N repeats and other apparent insertions, and disassembled/realigned. Phred(>Ewing et al. 1998) quality scores < 30 were investigated visually, and recoded as ambiguities (N) if not readily classified.

**Table 2.**
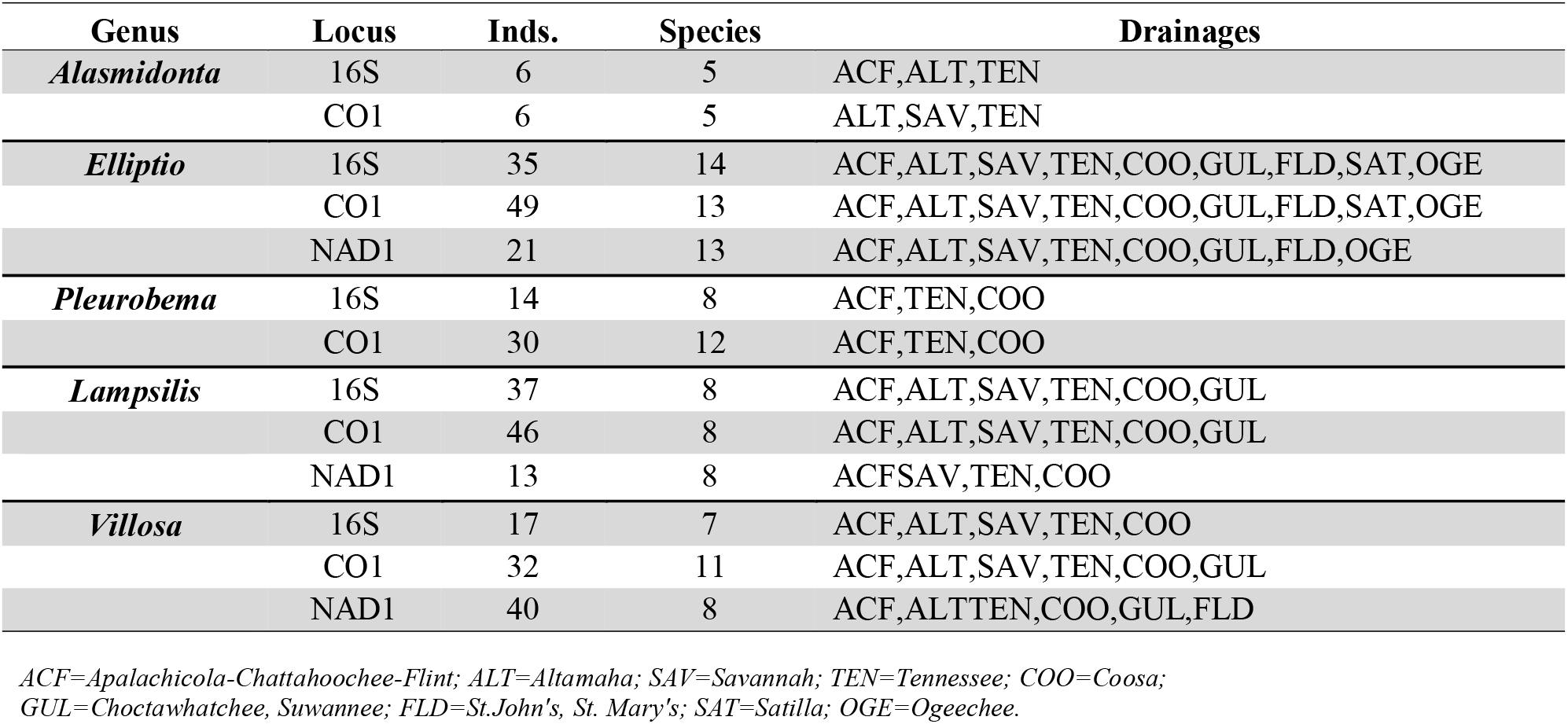
Summary of genetic data collected for freshwater mussels used in comparison among drainages.

### Phylogenetic Analysis

The program Beastversion 1.4.8 (Drummond and Rambaut 2007) was used to make inferences about demographic history based on mtDNA sequence phylogenies within the study populations. Beastis a Bayesian phylogenetic program that uses MCMC to sample the posterior distribution of gene trees, coalescence events and demographic parameters through time given observed DNA sequence data in addition to priors on substitution model and demographic models (Drummond and Rambaut 2007). All trees were rooted with the outgroup *Margaritifera margaritifera*, a freshwater mussel outside of the family Unionidae.

We used the program jModeltest (Posada 2008) to estimate which nucleotide substitution models best fitted the observed data. Priors on parameters associated with the substitution model in Beast were based on the values chosen by the Akaike information criterion (AIC) (Akaike 1974) in jModeltest. The best-fit model was HKY85+ Γ. For the molecular clock, we used a normal distributed substitution rate prior for *16S* with 95% of the probability density between 5.4×10–9 and 1.36×10–8, *CO1* and *NAD1* with a 95% of the probability density between 2.4×10–8 and 4.8×10–8 (Rawson and Hillbish 1995).

We used a relaxed molecular clock model as implemented in Beast, as preliminary analyses showed evidence of rate heterogeneity (Drummond, Ho et al. 2006). Species trees in Beast were estimated from aligned sequence data with enforcement of intraspecific monophyly and starting trees were randomly configured from a Yule prior (Aldous 2001), which assumes a constant speciation rate per lineage. All other parameters within Beast were given uniform distributions, where parameters were adjusted using the suggestions from the Beast log file (Drummond and Rambaut 2007). MCMC chains were run with 10^7^ iterations with trees sampled every 1000 iterations. The first 10% of the iterations were discarded as burn-in throughout. Log-files were analyzed in tracer version 1.4 (Rambaut &Drummond 2003), and effective sample sizes (ESS) were used to evaluate MCMC convergence within chains. logcombiner version 1.4.8 (Drummond and Rambaut 2007) was used to combine data from independent chains with identical settings into a composite chain to assess the robustness of parameter estimates.

## Results

We analyzed 346 sequences from five genera encompassing 37 species. Species were sampled across 12 distinct drainages all containing the Atlantic slope-Gulf of Mexico drainage split except *Pleurobema* (Table 2).

All Beast runs were found to have good convergence across 3 independent chains with an ESS of greater than 200. Mean substitution rates were found to be within reasonable bounds for both mitochondrial loci in invertebrates (Drake, Charlesworth et al. 1998). Drainage split times were inferred from the mean posterior distribution averaged over all loci within the sample. The mean substitution rate at each locus was used to estimate a divergence time between drainages and across the Appalachian Divide.

Gene phylogenies for each genus gave different topologies but “similar” chronologies. Overall the topologies of the different genera returned phylogenetic trees that were congruent with geographically contiguous drainages. The most common pattern was the grouping between the Atlantic slope and Gulf Coast drainages, however the Tennessee drainage was often the exception to this pattern grouping with the Atlantic Slope (Figure 2).

**Figure 2.**
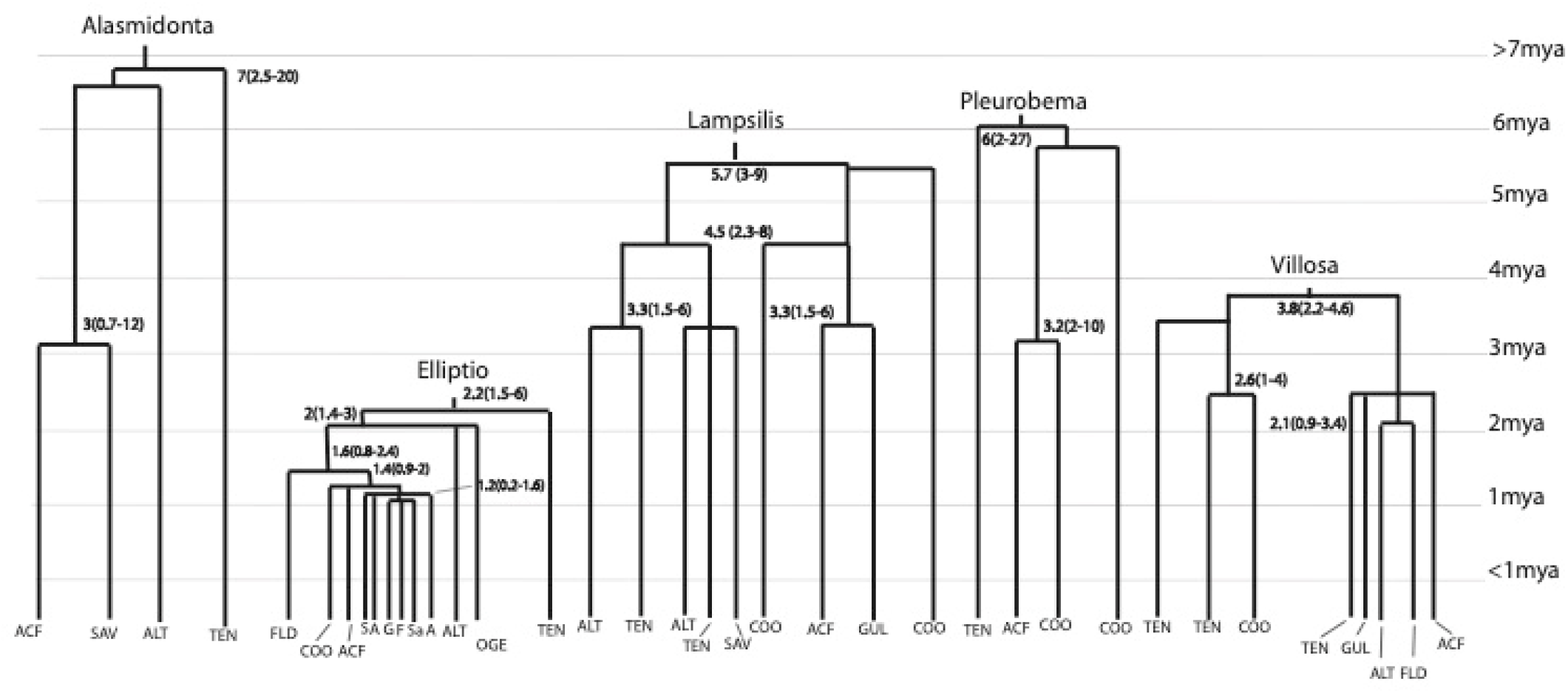
Phylogeny of five genera with divergence times by drainage. Comparison of species trees from five genera with tips labeled as river drainages. Mean age with 95% credible intervals on branching events are given using the mean substitution rate as given by Beast (Drummond &Rambaut 2007).

In the genus *Alasmidonta*, species from different drainages were reciprocally monophyletic with the Savannah and ACF having the fewest substitutions separating the terminal nodes. However, sampling issues limited us to comparing only the Savannah, Tennessee, and Altamaha, and thus restricting our ability to reconstruct drainages patterns using *Alasmidonta*. The genus *Elliptio* showed patterns of polyphyly in the Altamaha drainage with *Elliptio shepardiana* grouping with species from the Ogeechee drainage. Most other drainages for the genus *Elliptio* were reciprocally monophyletic, except Florida where limited sample size (n=2) does not allow us to make any concrete conclusions as to its placement. The genus *Lampsilis* also produced a polyphyletic Altamaha drainage with some species being more closely related to species from the Tennessee drainage than other Altamaha species. Other drainages within the *Lampsilis* genus were reciprocally monophyletic in respect to the Gulf Coast and Atlantic slope groupings with the exception of the Tennessee. Sparse sampling in the genus *Pleurobema* returned a polyphyletic Coosa drainage with a monophyletic Tennessee. The genus *Villosa* showed patterns of polyphyly in the Tennessee drainage with some species grouping with the Gulf drainages and some with the Coosa drainage, all other drainages were monophyletic.

Even though we found different topologies among the genera the chronologies were similar. Timing of speciation events were similar in *Alasmidonta, Lampsilis*, and *Pleurobema* at ∼5-7 millions of years ago (mya) with a more recent event affecting these genera as well as *Villosa* at ∼3 mya (Figure 2). The genera of *Elliptio* and *Villosa* share a speciation event between ∼2-3 mya with *Elliptio* having a more recent event at ∼1-2 mya (Figure 2).

### Appalachian Divide: Atlantic Slope-Gulf Coast split

We found evidence for two substantially different vicariance times corresponding to the Pliocene (*Alasmidonta*, and *Lampsilis*) and the Pleistocene (*Elliptio, Villosa*) across the Appalachian divide (Table 3). *Elliptio* species showed slightly different patterns across loci with an Atlantic-Gulf Coast split of 2.67 (1.7-3.7) mya using CO1 and earlier dates were found for *16S* 4.12 (0.6- 7) mya and *NAD1* 5.24 (2.3-9.0) mya owing mainly to the lack informative sites at *16S* and poor sampling for *NAD1. Villosa* species had a mean divergence time of 1.98 (1-1.3) mya for *CO1*, with earlier times for *16S* of 2.16 mya (1.2-4.2) and *NAD1* of 3.38 (2.2-4.6) mya. *Lampsilis* and *Alasmidonta* species showed evidence of an earlier split time, when compared to *Elliptio* and *Villosa* (Table 3).

**Table 3.** Summary of divergence times. Summary of divergence times for the Atlantic Slope-Gulf Coast estimated from mtDNA (*16S, CO1*, and *NAD1*) for each genus of mussel. We report the mean, median, and 95% credible intervals from the Bayesian analysis by Beast (Drummond &Rambaut 2007). Mean *μ* is the average rate of substitution per million years per gene averaged across the phylogeny. All phylogenies were rooted using the outgroup *Margaritifera margaritifera*.

| Genus | Locus | Mean (MYA) | Median (MYA) | 95 % CI | mean $\mu$ per million years |
| --- | --- | --- | --- | --- | --- |
| *Alasmidonta* | 16S | 12.83 | 11.35 | 4.2-25.3 | 0.010 |
|  | COI | 6.36 | 5.82 | 2.4-11.5 | 0.033 |
| *Elliptio* | 16S | 4.12 | 3.56 | 0.6-7 | 0.013 |
|  | COI | 2.67 | 2.59 | 1.7-3.7 | 0.036 |
|  | NAD1 | 5.24 | 4.90 | 2.3-9 | 0.042 |
| *Lampsilis* | 16S | 5.81 | 5.50 | 3-9.2 | 0.013 |
|  | COI | 3.92 | 1.45 | 0.3-14.7 | 0.036 |
|  | NAD1 | 5.62 | 5.38 | 2.3-9.9 | 0.042 |
| *Pleurobema* | 16S | 12.58 | 10.42 | 2.9-27.5 | 0.012 |
|  | COI | 6.93 | 6.52 | 3.4-11.3 | 0.037 |
| *Villosa* | 16S | 2.16 | 1.89 | 1.2-4.2 | 0.012 |
|  | COI | 1.98 | 1.90 | 1.1-3 | 0.037 |
|  | NAD1 | 3.38 | 3.30 | 2.2-4.6 | 0.052 |

Bayesian posterior estimates of divergence times per locus present a pattern of two separate vicariance events, one corresponding to *Elliptio* and *Villosa* at approximately 0.8-2.2 mya and a second corresponding to *Lampsilis* and *Alasmidonta* at approximately 4.8-6.5 mya. The 95% credible intervals for the posterior distribution between the *Elliptio-Villosa* group and the *Alasmidonta-Lampsilis* group do not overlap (Figure 3), further supporting two separate events.

**Figure 3.**
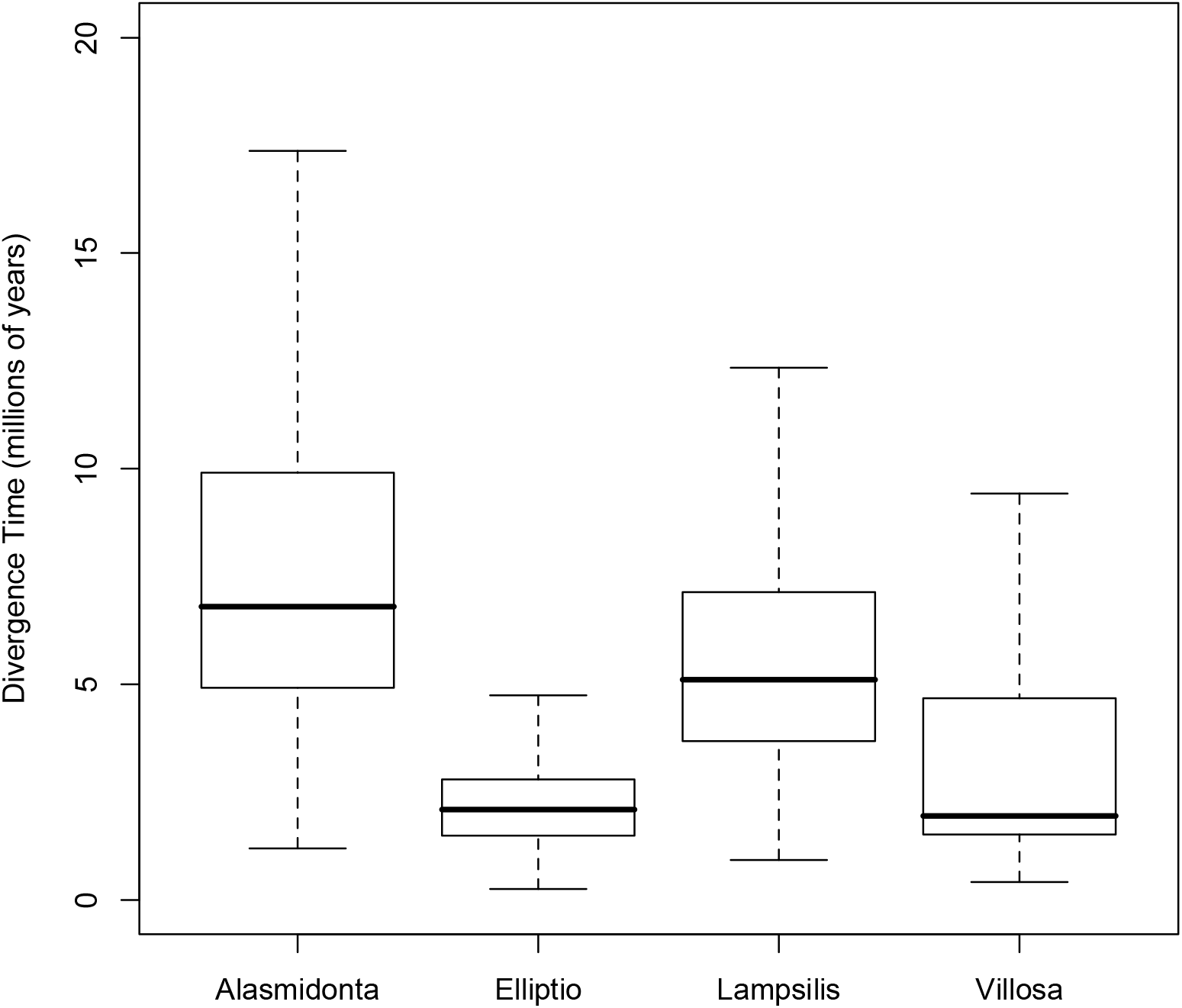
Divergence time between Atlantic and Gulf Coast drainages. Boxplot showing the divergence time between the Atlantic Slope and Gulf Coast drainages for four genera of freshwater mussels as average across all loci. Boxes delineate upper and lower quartiles, dark lines show medians, and dashed lines extend to the last observation within 1.5x interquartile range of the boxes.

## Discussion

In the southeast United States two main hypotheses have been proposed to explain the distribution of freshwater taxa among drainages. One proposed hypothesis is the source-pool where a central river high in species diversity seeded the surrounding rivers (Sepkoski and Rex 1974). This hypothesis proposed that various headwater captures, and flooding in the Pleistocene allowed the movement of species among adjunct rivers creating the present pattern of species distribution. A competing hypothesis proposed that fragmentation of a widespread ancestral species, existing before the Pleistocene, created the present pattern of species distribution. Mayden (1988) as well as others (Pflieger 1971; Mayden 1985; Mayden 1987; Wiley and Mayden 1985; Bermingham and Avise 1986; Avise 1992) provide support for a Pliocene vicariant event in freshwater fish taxa, refuting the source-dispersal hypothesis as the main mechanism responsible for contemporary freshwater fish species distributions. The current distribution of freshwater mussel species in the southeastern U.S. supports both the source- dispersal and vicariant hypotheses.

The hypotheses of source-dispersal and vicariance may not be mutually exclusive. This is in fact the case for freshwater mussels where we find evidence supporting both vicariance events and dispersal events. Vicariance events can be most clearly seen as the congruent bifurcation of phylogenetic trees among different genera. This pattern is evident in Figure 2 where the genera of *Lampsilis, Pleurobema*, and *Alasmidonta* share a diversification event and the genera of *Elliptio* and *Villosa* also share a diversification event. Earlier diversification events are also shared among genera such as the ∼3-4 mya diversification of *Lampsilis, Pleurobema, Alasmidonta*, and *Villosa* and the much earlier diversification event of *Villosa* and *Elliptio*. Congruence among diversification events is difficult due to confounding factors such as species life history, substitution rates, and ancestral population sizes (Edwards and Beerli 2000), but the support of multiple genera all with a similar pattern point to a general vicariant event that influenced the whole southeastern mussel fauna.

Dispersal events are identified by a species from a specific area, i.e. Coosa drainage, finding a most recent common ancestor with species from another drainage. Inferred dispersal events are rarer in the dataset possibly because of the exclusion of widespread species from the analysis. A few cases of dispersal are evident in the genera of *Lampsilis, Villosa*, and *Pleurobema* typically involving the Coosa or the Altamaha and Tennessee.

We find evidence for multiple divergence times in our five genera of mussels with *Elliptio* and *Villosa* species corresponding to a Pleistocene divergence time, while other genera had an earlier divergence time. Multiple divergence times were also found by Bermingham and Avise (1986) in the study of four fish species of the southeastern U.S that spanned the Appalachian divide. Bermingham and Avise (1986) found two separate divergence times, one for *Lepomis* dating to the Pliocene and a second later time for *Amia* dating to the Pleistocene. Bermingham and Avise (1986) propose that if a second vicariance event occurred in the Pleistocene it might have erased the earlier event in the genus *Amia*, while at the same time not affecting *Lepomis*. Pleistocene stream remodeling, notably the headwater captures of the Chattahoochee by the Savannah (Swift, Gilbert et al. 1986), may also have led to species exchange, but it may not be expected due to the rarity of many fish and mussel species in headwater reaches. In the species of *Elliptio* and *Villosa* we did not find any evidence supporting the movement of species via the headwater capture, but did find evidence in *Villosa* species that supports the Pleistocene connection of the Tennessee with the Mobile drainage through the presence of the hypothesized Appalachian River (Swift, Gilbert et al. 1986; Mayden 1988).

### Appalachian Divide: Atlantic-Gulf split

Our results provide evidence for a widespread pre-Pleistocene ancestor in species of *Alasmidonta* and *Lampsilis* with a deep phylogenetic divergence between Atlantic slope and Gulf Coast drainages of approximately 5-7 mya. However, we also found support for a more recent divergence in *Elliptio* and *Villosa* species dating to 0.7-1.8 mya also between Atlantic slope and Gulf Coast drainages.

Our estimated divergence times for the Atlantic-Gulf split agree with two other studies estimating vicariance in fish species of the Atlantic and Gulf Coast. Bermingham and Avise (1986) found a divergence time in *Lepomis* separating species of the Atlantic Slope drainages from the Gulf Coast drainages dating back to the Pliocene, and Mayden (1988) found evidence of a pre-Pleistocene vicariance event in seven fish taxa of the Eastern and Central highlands. Bermingham and Avise (1986) offer the explanation that saltwater inundation of the lowlands during the Pliocene interglacial (50-80m above present-day sea levels), isolated and fragmented a once widespread species. Then as the seas receded from the Pliocene high sea-level stand, it would have been possible for fish to disperse along the lowlands, enabling taxa within major lineages to colonize adjacent coastal rivers.

The major genetic effects of the Pliocene 1 million year-long, high sea-level stand on both mussels and fish species would have been *i)* extinction of locally differentiated species in the smaller Coastal Plain rivers, *ii)* attendant reduction of overall levels of genetic diversity within each species, *iii)* significant sequence divergence between lineages that had survived in refugia of either piedmont headwaters or Floridian highlands (Bermingham and Avise 1986; Baer 1998). Isolation of mussel taxa with fish taxa for long periods of time would also have allowed for the specification of host-parasite relationships and may account for some of the contemporary patterns of species-specific parasitism see among different genera of mussels, where some mussel species are specialists and other are generalists.

### Reproductive isolation in mussels

Species formation and reproductive isolation are poorly understood in freshwater mussel taxa. It has been suggested that mechanisms such as lysin recognition proteins similar to the genus *Mytilus* may be acting to prevent species hybridization (Riginos and McDonald 2003), but there is also the possibility of reproductive isolation via a breakdown of fish-host relationships with hybrid individuals (Kat 1985; Kat 1986). If vicariance events are responsible for most species’ formation then the presence of multiple endemic species within drainage may be due to sympatric speciation rather than reticulate allopatry (van Veller, Kornet et al. 2000).

Broadening our understanding of vicariance events and their frequency between drainages serve to inform future hypotheses of speciation and reproductive isolation in freshwater mussel species. If our results on mussel vicariance, as well as the corroboration of data from fish taxa, are any indication of speciation processes in the Unionids it might be possible that while the first burst of species formation arose via allopatric speciation, while subsequent, within drainage endemism is due entirely to sympatric of parapatric speciation. This hypothesis arises from the strong reciprocal monophyly found in most mussel and fish species for a particular drainage. As most fish and mussel species find a most recent common ancestor within a drainage before finding an ancestor between drainages, it could be assumed that after the initial allopatric separation, a secondary burst of species formation occurred within drainages. However, our data suggests that within-drainage speciation may have occurred as recently as 1 mya in species of *Lampsilis* in the Coosa drainage. Species within drainages are typically reciprocally monophyletic but harbor vast amounts of genetic diversity leading to shallow branches between species. Speciation within a drainage either via sympatric or parapatric speciation would most likely require the formation of prezygotic barriers to reproductive isolation more so than in the case of allopatric speciation (Coyne and Orr 2004). However, it is possible that within drainage speciation events are linked to introductions of new fish hosts by dispersal from other drainages, which would allow specialization and possible diversification into different species.

## Conclusions

When two or more groups display patterns congruent in time and space, the patterns are probably the result of common history. Our paper tests the concordance of species divergence with glacial and interglacial vicariance to determine whether divergence times are temporally congruent among mussel genera. Our ability to differentiate between hypotheses that have significantly influenced the ecological and evolutionary history of the biota in the southeast United States will assist in taxonomic reconstruction of species relationships within the region and allow inference into long-term community stability in freshwater ecosystems.

## Acknowledgements

The authors would like to thank A. Bogan, D. Campbell, M. Gangloff, J. Smith, for help with procuring museum collections; M. Raley, J.Wisniewski, for identification of endemic species and recent nomenclature changes; and the Wares Lab members (2007-2009) for discussion.

## References

Akaike, H. (1974). “A new look at the statistical model identification”. IEEE Transactions on Automatic Control 19 (6): 716–723.

Aldous D.J. (2001). “Stochastic models and descriptive statistics for phylogenetic trees, from Yule to today.” Statistical Science 16(1):23–34.

Avise, J.C. (1992). “Molecular population structure and the biogeographic history of a regional fauna: a case history with lessons for conservation biology.” Oikos 63:62–76.

Avise, J.C. (1998). “The history and purview of phylogeography: a personal reflection.” Molecular Ecology 7:371–379.

Baer, C. F. (1998). “Species-Wide population structure in a southeastern U.S. freshwater fish, Heterandria Formosa: geneflow and biogeography.” Evolution 52: 183–193

Bermingham, E. and J.C. Avise. (1986). “ Molecular zoogeography of freshwater fishes in the southeastern United States.” Genetics 113: 939–965

Bermingham, E., and C. Moritz. (1998). “Comparative phylogeography: concepts and applications.” Mol. Ecology 7: 367–369.

Campbell, D. C., J. M. Serb, et al. (2005). “Phylogeny of North American amblemines (Bivalvia, Unionoida): prodigious polyphyly proves pervasive across genera.” Invertebrate Biology 124(2): 131–164.

Coyne, J.A. and H.A. Orr. (2004). Speciation. Sinauer Associates, Sunderland, MA.

Drake JW, Charlesworth B, et al. (1998). “Rates of spontaneous mutation.” Genetics 148: 1667–1686.

Drummond A.J., S.Y.W. Ho, et al. (2006). “ Relaxed phylogenetics and dating with confidence.” PLoS Biology 4(5)

Drummond, A.J. and A. Rambaut. (2007). “beast: Bayesian evolutionary analysis by sampling trees.” BMC Evolutionary Biology, 7, 214.

Edwards, S.V. and P. Beerli. (2000). “Gene divergence, population divergence, and the variance in coalescence time in phylogeographic studies.” Evolution 54(6): 1839–1854.

Folmer, O., M. Black, et al. (1994). “DNA primers for amplification of mitochondrial cytochrome c oxidase subunit I from diverse metazoan invertebrates.” Molecular Marine Biology and Biotechnology 3: 294–299.

Gilbert, C. R. (1987). “Zoogeography of the freshwater fish fauna of southern Georgia and peninsular Florida.” Brimleyana 13:25–54.

Grande, L. (1990). “Vicariance biogeography” in Paleobiology, a synthesis. (ed. By D.E.G. Briggs & P.R. Crowther), pp. 448–451. Blackwell Scientific Publications, Oxford.

Jarman, S., D. W. Robert, et al. (2002). “Oligonucleotide Primers for PCR Amplification of Coelomate Introns.” Marine Biotechnology 4(4): 347–355.

Johnson, R.I. (1970). “The systematics and zoogeography of the Unionidae (Mollusca: Bivalvia) of the Southern Atlantic Slope region.” Bull Mus. Comp Zool. 140:263–450.

Johnson, R.I. (1972). “The Unionidae (Mollusca: Bivalvia) of Peninsular Florida.” Bull. Florida St. Mus. 16:181–249.

Kat, P. W. (1985). “Historical evidence for fluctuation in levels of hybridization.” Evolution 39:1164–1169.

Kat, P.W. (1986). “Hybridization in a unionid faunal suture zone.” Malacologia 27:107–125.

Lieberman, B.S. (2000). “Applying molecular phylogeography to test paleoecological hypotheses: a case study involving Amblema plicata (Mollusca, Unionidae).” Pp. 83-103 in W. D. Allmon and D. Bottjer (eds.), Evolutionary Paleoecology. Columbia University Press, New York.

Martin, F.D. (1980). “Heterandria formosa Agassiz, least killifish.” P. 547 in D. S. Lee, C. R. Gilbert, C. H. Hocutt, R. E. Jenkins, D. E. McAllister, and J. R. Stauffer Jr., eds. Atlas of North American fishes. North Carolina State Museum of Natural History, Raleigh.

Master, L. (1990). “The imperiled status of North American aquatic animals.” Biodiversity Network News 3:1–2.

Mayden, R.L. (1988). “Vicariance biogeography, parsimony, and evolution in North Amercian Freshwater fishes.” Systematic Zoology 37: 329–355.

Mayden, R.L. (1985). “Biogeography of Ouachita Highland fishes.” Southwest. Nat., 30p:195-211.

Mayden, R.L. (1987a). “Historical ecology and North American Highland fishes: A research program in community ecology.” Pages 210-222 in Community and evolutionary ecology of North American stream fishes (W. J. Matthews and D. C. Heins, eds.). University of Oklahoma Press, Norman.

Mayden, R.L. (1987b). “Pleistocene glaciation and historical biogeography of North American central highland fishes.” Pages 141-151 in Quaternary environments of Kansas (W. C. Johnson, ed.). Kansas Geological Survey, Guidebook Series 5.

Page, R.D.M. (1994). “Parallel phylogenies: reconstructing the history of host-parasite assemblages.” Cladistics 10: 155–173.

Pflieger, K. (1971). A distributional study of Missouri fishes. Univ. Kansas, Mus. Nat. Hist., Publication Series, 20(3):226–570.

Posada, D. (2008). “jModelTest: Phylogenetic Model Averaging.” Molecular Biology and Evolution 25: 1253–1256.

Rambaut, A. and A.J. Drummond (2003). “Tracer [computer program]”Available from http://evolve.zoo.ox.ac.uk/software/2003

Rawson, P.D. and T. J. Hilbish. (1995). “Distribution of male and female transmitted mitochondrial lineages in the mussels Mytilus trossulus and M. galloprovincialis on the west coast of North America.” Mar. Biol. 124:245–250.

Riginos, C. and J.H. McDonald. (2003). “Positive selection on an acrosomal sperm protein, M7 Lysin, in three species of Mussel Genus Mytilus.” Molecular Biology and Evolution 20:200–207.

Rozas, J., J.C. Sanchez-DelBarrio, et al. (2003). “DnaSP, DNA polymorphism analyses by the coalescent and other methods.” Bioinformatics 19: 2496–2497.

Sepkoski, J. J. and M.A. Rex. (1974). “Distribution of freshwater mussels: coastal rivers as biogeographic islands.” Systematic Zoology 23: 165–188.

Swift, C.C., C.R. Gilbert, et al. (1986). “Zoogeography of the freshwater fishes of the southeastern United States: Savannah River to Lake Ponchartrain.” Pp. 213–266 in C. H. Hocutt and E. Wiley, eds.

van Veller, M.G.P., D.J. Kornet, et al. (2000). “Methods in Vicariance biogeography: Assesment of the implementations of assumptions 0,1, and 2.” Cladistics 16: 319–345.

Wares JP, Pankey MS, et al. (2009). “A “shallow phylogeny” of shallow barnacles (Chthamalus).” PLoS One in press.

Wiley, E. O. and R.L. Mayden. (1985) “Examples from the North American fish fauna.” Ann. M. Bot. Card. 72: 596–635.

Williams, J.D., A.E. Bogan, et al. (2008). Freshwater Mussels of Alabama and the Mobile Basin in Georgia, Mississippi, and Tennessee. University Alabama Press, AL.

Wooten, M.C., K. T. Scribner, et al. (1988). “Genetic variability and systematics of Gambusia in the southeastern United States.” Copeia 1988:283–289.

